# Variable responses of individual species to tropical forest degradation

**DOI:** 10.1101/2024.02.09.576668

**Authors:** Robert M. Ewers, William D. Pearse, C. David L. Orme, Priyanga Amarasekare, Tijmen De Lorm, Natasha Granville, Rahayu Adzhar, David C. Aldridge, Marc Ancrenaz, Georgina Atton, Holly Barclay, Maxwell V. L. Barclay, Henry Bernard, Jake E. Bicknell, Tom R. Bishop, Joshua Blackman, Sabine Both, Michael J. W. Boyle, Hayley Brant, Ella Brasington, David F.R.P. Burslem, Emma R. Bush, Kerry Calloway, Chris Carbone, Lauren Cator, Philip M. Chapman, Vun Khen Chey, Arthur Chung, Elizabeth L. Clare, Jeremy Cusack, Martin Dančák, Zoe G. Davies, Charles W. Davison, Mahadimenakbar M. Dawood, Nicolas J. Deere, Katharine J. M. Dickinson, Raphael K. Didham, Timm F. Döbert, Rory A. Dow, Rosie Drinkwater, David P. Edwards, Paul Eggleton, Aisyah Faruk, Tom M. Fayle, Arman Hadi Fikri, Robert J. Fletcher, Hollie Folkard-Tapp, William A. Foster, Adam Fraser, Richard Gill, Ross E. J. Gray, Ryan Gray, Nichar Gregory, Jane Hardwick, Martina F. Harianja, Jessica K. Haysom, David R. Hemprich-Bennett, Sui Peng Heon, Michal Hroneš, Evyen W. Jebrail, Nick Jones, Palasiah Jotan, Victoria A. Kemp, Lois Kinneen, Roger Kitching, Oliver Konopik, Boon Hee Kueh, Isolde Lane-Shaw, Owen T. Lewis, Sarah H. Luke, Emma Mackintosh, Catherine S. Maclean, Noreen Majalap, Yadvinder Malhi, Stephanie Martin, Michael Massam, Radim Matula, Sarah Maunsell, Amelia R. Mckinlay, Simon Mitchell, Katherine E. Mullin, Reuben Nilus, Ciar D. Noble, Jonathan M. Parrett, Marion Pfeifer, Annabel Pianzin, Lorenzo Picinali, Rajeev Pillay, Frederica Poznansky, Aaron Prairie, Lan Qie, Homathevi Rahman, Terhi Riutta, Stephen J. Rossiter, J. Marcus Rowcliffe, Gabrielle Briana Roxby, Dave J. I. Seaman, Sarab S. Sethi, Adi Shabrani, Adam Sharp, Eleanor M. Slade, Jani Sleutel, Nigel Stork, Matthew Struebig, Martin Svátek, Tom Swinfield, Heok Hui Tan, Yit Arn Teh, Jack Thorley, Edgar C. Turner, Joshua P. Twining, Maisie Vollans, Oliver Wearn, Bruce L. Webber, Fabienne Wiederkehr, Clare L Wilkinson, Joseph Williamson, Anna Wong, Darren C. J. Yeo, Natalie Yoh, Kalsum M. Yusah, Genevieve Yvon-Durocher, Nursyamin Zulkifli, Olivia Daniel, Glen Reynolds, Cristina Banks-Leite

## Abstract

The functional stability of ecosystems depends greatly on interspecific differences in responses to environmental perturbation. However, responses to perturbation are not necessarily invariant among populations of the same species, so intraspecific variation in responses might also contribute. Such inter-population response diversity has recently been shown to occur spatially across species ranges, but we lack estimates of the extent to which individual populations across an entire community might have perturbation responses that vary through time. We assess this using 524 taxa that have been repeatedly surveyed for the effects of tropical forest logging at a focal landscape in Sabah, Malaysia. Just 39 % of taxa – all with non-significant responses to forest degradation – had invariant responses. All other taxa (61 %) showed significantly different responses to the same forest degradation gradient across surveys, with 6 % of taxa responding to forest degradation in opposite directions across multiple surveys. Individual surveys had low power (< 80 %) to determine the correct direction of response to forest degradation for one-fifth of all taxa. Recurrent rounds of logging disturbance increased the prevalence of intra-population response diversity, while uncontrollable environmental variation and/or turnover of intraspecific phenotypes generated variable responses in at least 44 % of taxa. Our results show that the responses of individual species to local environmental perturbations are remarkably flexible, likely providing an unrealised boost to the stability of disturbed habitats such as logged tropical forests.

## Introduction

Species differ in their traits, with the implication that they will respond differently to the same environmental perturbation (1, 2). This interspecific “response diversity” has been identified as a key determinant of community and ecosystem stability for several decades (3, 4). Yet there is newly emerging evidence that perturbation responses within species are surprisingly variable, because any given species might respond differently to the same perturbation depending on where it experiences that perturbation (5): the population-level responses of individual species vary according to position within their geographic range (6), climate envelope (7) and macroclimatic conditions (8).

Consequently, the “response traits” that purportedly define how a given species will respond to environmental change (9) may not have a fixed relationship with species’ morphological traits (10), despite this being widely assumed in many analyses (2, 11, 12). What we don’t yet know, however, is whether the population-level responses at a single location are fixed and invariant through time, or whether those responses might also vary. Such variation might arise through population-level turnover in the phenotypes of individuals, and in response to the ever-changing environmental conditions in which local ecosystems are embedded. If it exists, any such local, intra-population response diversity through time might be expected to boost the stability of disturbed ecosystems (3, 13).

Here, we directly quantified the degree of intra-population response diversity by comparing taxon-specific occurrence patterns described in multiple surveys that were collected within a single landscape: the SAFE Project in Sabah, Malaysia (14). Data were collected along a forest degradation gradient defined by variation in logging intensity, which we quantify as the percentage of biomass reduction. Logging in tropical rainforests is often a recurrent perturbation, and our sites are no different. Sites were logged between zero and four times between 1970 and 2008 (15), and many underwent an additional round of salvage logging between 2012 and 2015. Sites at SAFE therefore extend across all levels of biomass reduction, encompassing primary forest with no (0 %) biomass removal, areas of light and moderate selective harvesting of trees, through to salvage logged and clear-felled sites with virtually all (100 %) above-ground biomass removed. Habitat degradation gradients of this magnitude should generate strong, predictable impacts on the occurrence patterns of individual taxa, and as such it represents a good system in which to quantify the degree of intra-population response diversity.

Our analysis encompassed 119 single-year surveys that included at least one taxon that was also sampled in at least one other survey. Individual surveys – including those conducted on the same taxon – varied in one or more dimensions of survey method, sample sites and year (Table S1), reflecting variation in the study design process of individual researchers. While all surveys were conducted within a single year, we stress that our definition of single-year survey does not imply that each survey conducted just a single site visit. Sixty-six of the 119 surveys (55 %) conducted repeat site visits within the year, meaning the researchers had sampled more intensively than a snapshot survey in which a site is visited just a single time. Data from multiple site visits within a year were aggregated to represent a single survey for analysis.

There were 1,258 taxa observed in two or more of these spatially overlapping, single-year surveys, of which 524 had high enough occurrences to be modelled in multiple surveys (n ≥ 5 occurrences in each of ≥ 2 surveys). Sensitivity analysis demonstrated that this choice of occurrence threshold ensured the results and conclusions we present are conservative estimates of response diversity (SI Appendix; Fig. S1). The 524 taxa included 122 plants, 205 invertebrates, 17 fish, 2 reptiles, 17 amphibians, 100 birds and 61 mammals. We focus our analysis on patterns of taxon occurrence. Taxon occurrence is a simple, commonly employed analysis of biodiversity patterns that should be more robust to among-year variation in population sizes than analyses based on abundance (16, 17), and therefore represents a conservative test of response diversity. We fitted binomial general linear models to presence-absence data for 1,942 taxa × survey combinations, from which we recorded two metrics: (1) the statistical significance of the occurrence pattern, categorised into significant (*p* < 0.05) or non-significant (*p* ≥ 0.05) (sensitivity analysis demonstrated little impact of the choice of *p*-value on our conclusions; SI Appendix and Fig. S1); and (2) the slope and intercept of the occurrence pattern. These two metrics can be combined to define four types of response diversity in empirical data (18) (Fig. 1): (A) invariant, where responses are statistically indistinguishable; (B) magnitude, where all responses are statistically significant and have a common direction, but have variable slope estimates; (C) uncertainty, where some responses are statistically significant but others are not; and (D) sign changes, where statistically significant responses occur but in opposite directions.

**Figure 1.**
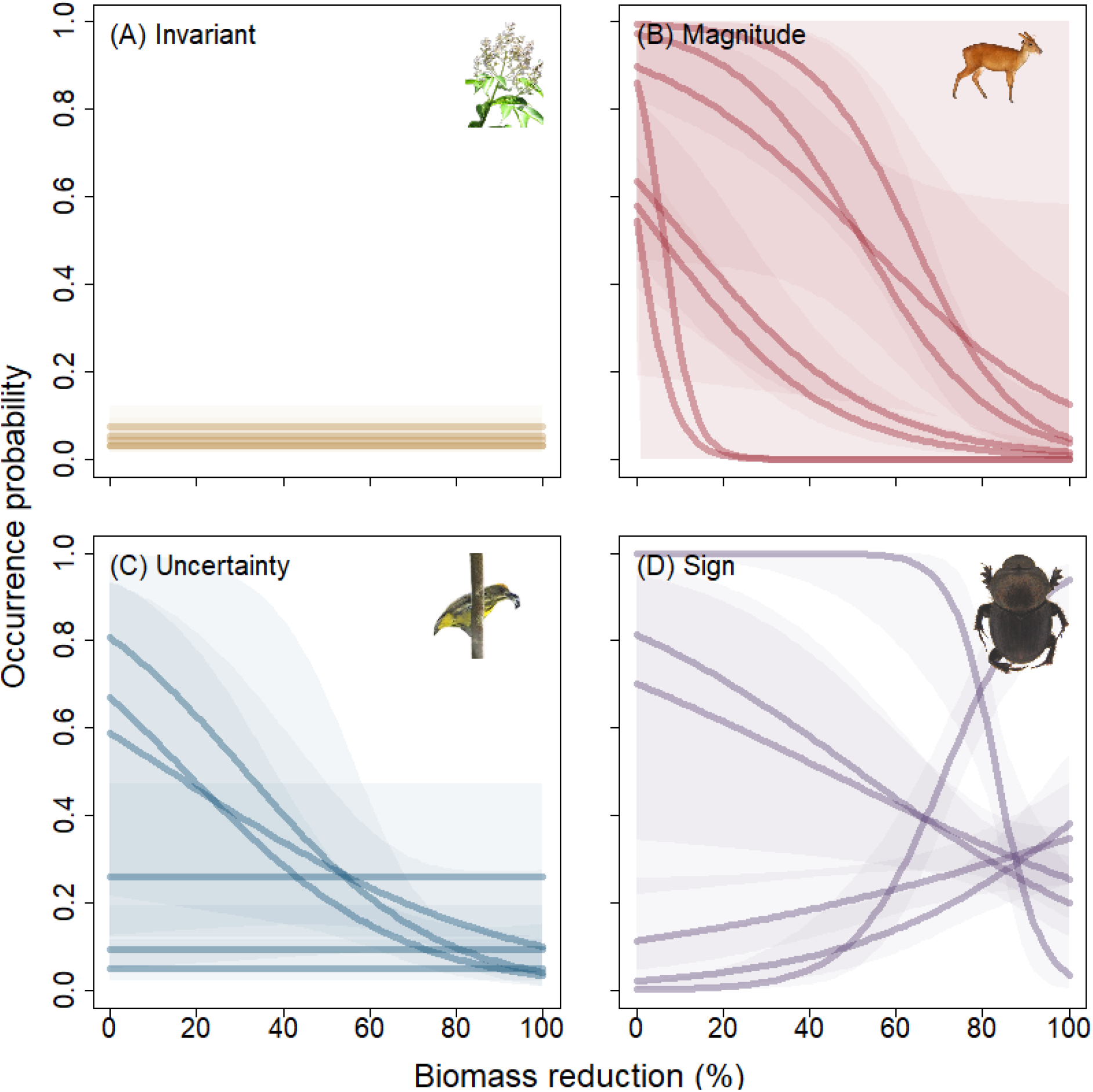
Examples of intra-population response diversity to a gradient of forest degradation. In all panels, forest degradation is represented as a percentage reduction in aboveground biomass, where zero represents the median biomass in unlogged forest. Each line displays a fitted model generated to empirical data collected from a different survey of that particular taxon. Shaded polygons represent the 95 % confidence intervals. Statistically non-significant relationships are displayed as an intercept-only fitted model. **(A)** Invariant: all observed responses of that taxon to forest degradation in different surveys are statistically indistinguishable (e.g. the tree genus Vitex). **(B)** Magnitude: responses of that taxon are all statistically significant and have the same direction of effect, but have slope and/or intercept estimates that differ (e.g. Bornean yellow muntjac Muntiacus atherodes). **(C)** Uncertainty: responses of that taxon vary in their statistical significance (e.g. Yellow breasted flowerpecker Prionochilus maculatus). **(D)** Sign: responses of that taxon are statistically significant, but have response patterns in opposing directions (e.g. dung beetle Onthophagus mulleri).

### High intra-population response diversity of tropical forest taxa

Almost two-thirds (61 %) of all taxa exhibited response diversity, and our consistently conservative approach to the analysis means this estimate represents a lower bound (Fig. S1). We found 154 taxa (29 %) with magnitude response diversity, having response patterns that were consistent in terms of statistical significance and directions of effect, but where the slope or intercept of the observed effect varied significantly among surveys (Fig. 1B). Most commonly, response diversity was in the form of uncertainty, (n = 254, 48 %; Fig. 2A), with some surveys finding non-significant response patterns but others finding statistically significant trends (Fig. 1C). This proportion is higher than would be expected by chance (12-41 %, SI Appendix), indicating this result is not a spurious one emerging from the combination of Type I (false positive) and Type II (false negative) sampling errors across multiple surveys. Finally, 6 % of taxa (n = 32) displayed the most extreme form of intra-population response diversity – sign diversity – where repeated surveys detected statistically significant response patterns in opposing directions (Fig. 1D). The latter three classes are not mutually exclusive, and 23 % of taxa (n = 122) exhibited multiple forms of response diversity (Fig. 2A). For example, a taxon with three surveys might have one with a non-significant result and two with statistically significant responses in opposite directions, demonstrating both uncertainty and sign class. Our results provide quantitative insight into the intra-population response diversity of individual taxa to an environmental gradient at a single location, and indicate that more than one half of tropical forest taxa might demonstrate remarkably flexible responses to habitat degradation.

**Figure 2.**
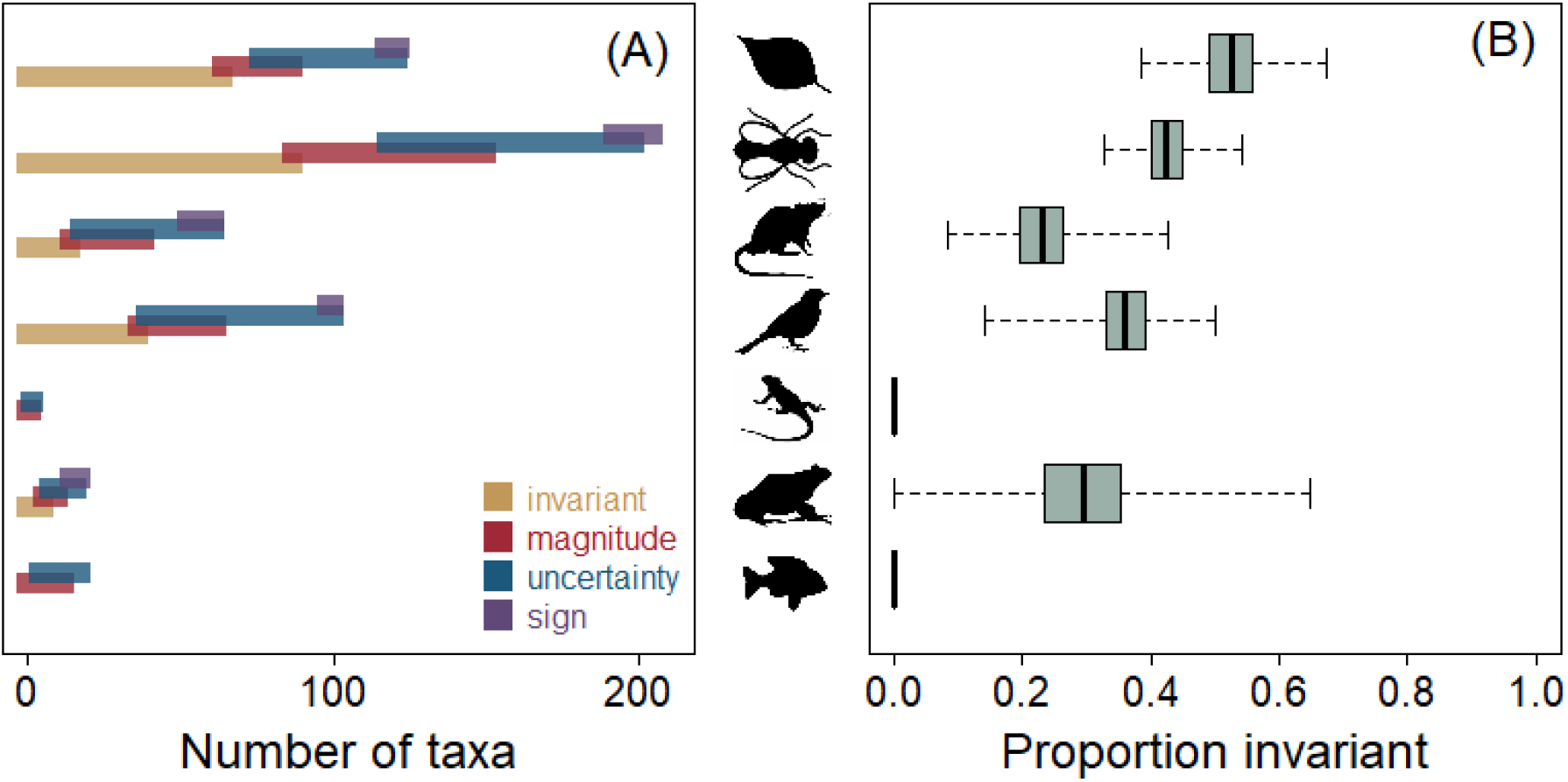
Broad taxonomic patterns in the replicability of single-year biodiversity surveys. **(A)** Number of taxa exhibiting each of the four classes of intra-population response diversity (see Fig. 1 caption for definitions). Classes are not all mutually exclusive, which is displayed with partially overlapping bars. **(B)** Bootstrapped estimates of the proportion of taxa with invariant, statistically indistinguishable response patterns across multiple surveys. Thick line represents the median, boxes the 1^st^ and 3^rd^ quantiles respectively, and whiskers the range.

Less than half of all taxa (39 %; n = 206) had invariant responses to the forest degradation gradient (Fig. 2A), in which all surveys gave results that were indistinguishable in terms of their statistical significance, direction and magnitude of effect (Fig. 1A). The proportion of taxa with fully invariant results varied across broad taxonomic groupings, varying from zero in fish to more than half for plants (Fig. 2B). However, all of the 206 taxa (100 %) with invariant response patterns also had no significant response to the forest degradation gradient in any survey. By contrast, there were 318 taxa that exhibited a significant response in at least one survey, and not one of those (0 %) responded in a fully invariant manner in all surveys, suggesting intra-population response diversity is the norm among taxa with meaningful habitat preferences.

Some of the intra-population response diversity we observed can be ascribed to life history characteristics of the taxa. The number of taxa exhibiting each of the four response diversity classes differed from a null expectation for all taxonomic groups (Fig. S2; *χ*^2^ Goodness of fit test, 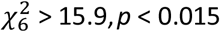). Plants were more likely to have invariant responses than expected by chance, reflecting the fact that trees are long-lived and stationary organisms. Repeated surveys of vegetation plots are always likely to detect the same individuals and thereby generate invariant response patterns. However, different studies with alternative spatial designs, or that examine different life history stages such as adults *versus* seedlings, still generated variable results. Mammals, by contrast, were more likely to exhibit response diversity in the form of both the magnitude and the sign classes. This pattern probably reflects the highly mobile nature of large mammals, which might allow them to rapidly re-distribute in response to continuously changing local conditions, including the spatial and temporal patchiness of fruiting events.

### Causes of intra-population response diversity

Intra-population response diversity can arise through spurious or ecologically meaningful mechanisms. First, variation in study design and field methods might generate spurious differences in observed response patterns that have nothing to do with the ecology of the species themselves. By contrast, ecologically meaningful response diversity might arise in one of two ways. First, variation in the phenotype of individuals can influence how they respond to disturbances. Any temporal turnover in the phenotypic composition of populations, which could be driven by selection forces linked to the forest degradation or varying environmental conditions, might contribute to the intra-population response diversity we have observed. We have no repeated morphological or genomic measurements that would allow us to detect or quantify this effect. And second, meaningful response diversity that might arise from uncontrollable and unmeasured environmental variation (18). Specific examples might include human disturbance such as hunting intensity and logging activity, and year-to-year variation in animal population sizes and movements driven by phenological events such as fruiting, or by climatic variation and extremes such as El Niño oscillations. When comparing the occurrence patterns from two surveys, we can only definitively rule out spurious response diversity, and by elimination confirm the presence of meaningful response diversity, if a pair of surveys used identical sampling methods in identical sampling sites. In these cases, any inconsistency in the taxon-specific response patterns between surveys must be due to either unaccounted for environmental variation or phenotypic turnover.

We confirmed the presence of meaningful response diversity in 44 % of taxa (174 out of 397 that had overlapping methods and sampling sites). A roughly equal number of taxa (n = 168, 42 %) had variable response patterns that could not be definitively confirmed as being meaningful due to variations in sampling sites and/or methods, and we therefore consider them to be confirmed examples of spurious response diversity. Both estimates should be considered to represent the lower bound of the actual values, however, as any survey pair could simultaneously exhibit both meaningful and spurious response diversity and there is no statistical method that can quantify this overlap.

We investigated three potential hypotheses that might explain the presence of meaningful intra-population response diversity in our data. There was no effect of the number of years between surveys on the probability of two surveys giving invariant results (Fig S3A; binomial GLM: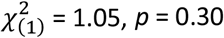), nor was there a discernible effect of El Niño events (Fig. S3B;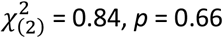), despite the latter having impacted forest growth patterns both through time and across space at our study site (19). We did, however, find that an extended logging event that occurred in the middle of our decade of observations, and that further reduced the biomass across large portions of the SAFE Project study area, influenced the pattern of survey results (Fig. S3C;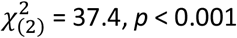). We found increased intra-population response diversity for survey pairs occurring within the logging events relative to survey pairs occurring in non-disturbed years, indicating that species responses are more variable during extreme land use change events. This increased response diversity did not occur equally across taxonomic groups (logging x taxonomic group interaction effect: *x*^2^ = 41.5, *p* < 0.001), but was instead driven by a reduction in the proportion of invariant responses of bird and mammal taxa. These taxa both have relatively high mobility, so their increased response diversity may reflect a tendency to move away from active disturbances.

Spurious response diversity, meanwhile, arose from variation in the methodology and fine details of individual studies, with the exact effect varying almost species-by-species (SI Appendix). One possible implication is that variation in the way sample sites are distributed along an environmental gradient may exert considerable, undetected influence over the outcome of a biodiversity survey.

Even unconscious bias in the choices of individual researchers about exactly where in the vicinity of a planned sample point to set a quadrat or place a trap could be affecting the outcome of surveys (20).

### Generating reliable results from ecological surveys

Much of ecology and conservation relies on single-year and snapshot surveys. Close to half (44 %) of the studies published in the journal *Ecology* present results based on data from a single year, and many of the world’s largest conservation NGOs rely on rapid biodiversity assessments to prioritise their actions (21). Drawing general inference from single-year studies on taxa with intra-population response diversity could easily lead to misleading conclusions about biodiversity patterns (17), and these could, in turn, lead to poor management and conservation decision-making.

We estimate that any given taxon needs to be analysed in each of three surveys to gain reliable insight into the impacts of forest degradation (Fig. 3A). Of the four response diversity classes, getting the direction (sign) of an effect wrong is the most immediately problematic: it could lead directly to management decisions that are the exact opposite of what is needed. We therefore used Bayesian hierarchical models to estimate the observed variation within and among taxon-specific surveys, and so determine the probability that analysis of a single-year survey will return the correct sign. This probability was less than 80 % – the assumed power of standard statistical tests – for one fifth of all taxa (n = 109; 21 %), but varied significantly among taxonomic groups (Fig. 3B, Kruskal-Wallis: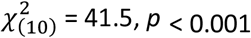). Birds were most likely to return correct signs from a single survey, while invertebrates were the least likely. On average, three surveys were needed to ensure a 90 % probability of getting the right direction of effect for 90 % of taxa (Fig. 3).

**Figure 3.**
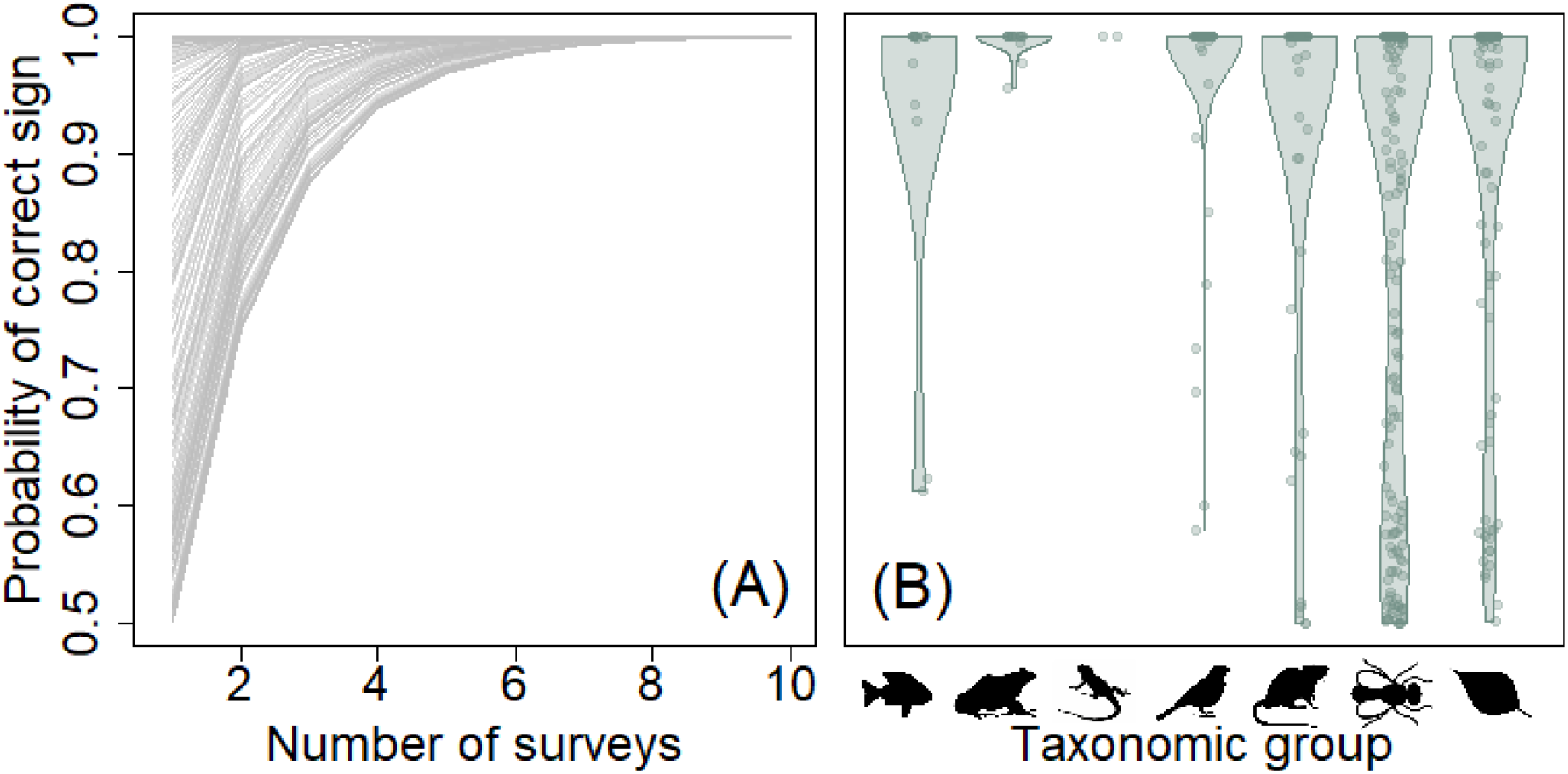
The probability of an analysis returning the correct sign (direction) of a taxon’s response to forest degradation as a function of the number of single-year surveys in which that taxon was detected. (A) Each grey line represents an analysis for one taxon (n = 494). (B) Violin plots showing the distribution of probabilities of a single-year survey returning the correct sign for seven taxonomic groups. Points indicate the probability for individual taxa within each group.

### Implications for ecology and conservation management

Our analyses raise the uncomfortable possibility that three out of every five taxon-specific results published in single-year studies may be unreliable representations of biodiversity patterns, whether that be through spurious methodological issues or due to meaningful intra-population response diversity. Yet our results should not be a complete surprise and do not appear to be unique to our study site. Inter-annual variation in the spatial distribution of species has been reported for taxa as diverse as birds in Australia (17), stream invertebrates in Finland (22) and plants in China (23). Similarly, it is becoming apparent that data spanning a decade or more are needed to obtain consistent results in ecological field experiments (24, 25), and time lags in the responses of species to environmental impacts (26) can mean the results of short-term studies can generate unrepresentative results.

Given such high levels of intra-population variation in response patterns, how can we generate conclusive insights in ecology and conservation? Intra-population response diversity means clear, definitive answers to biodiversity and conservation questions cannot be obtained from short funding cycles. Grants to collect new field data need to be awarded for longer durations, the continuation of multi-taxa biodiversity time series along environmental gradients should be prioritised, and studies specifically targeted at repeating earlier work should be supported. Until then, the most efficient way forward will likely be to find new ways that reliably transfer results among study sites. Early indications are that such transferability may be low (5), but for predictable reasons including the location of a site relative to species’ geographic ranges (6) and climatic tolerances (7). Clearer understanding of how these macroecological patterns influence site- and species-specific biodiversity patterns will be key to developing new predictive frameworks to extrapolate findings from heavily studied sites to regions lacking equivalent data (5). Such frameworks will provide a means to maximise the utility of the data that do exist, help contextualise the generality of ecological conclusions arising from single-year studies, and give the empirical evidence needed to support urgent decision making at national and regional scales.

The functional stability of ecosystems can be promoted through the diversity of relationships between species and the environment (3), with communities containing suites of species that vary in their responses to environmental gradients better able to stabilise ecosystem properties than communities lacking equivalent response diversity (4). Temporal or spatial variation in the responses of individual populations is similarly expected to contribute to the stability of ecosystems (27), but that contribution is unlikely to be equal among the different classes of response variability: we expect variation in the sign of a response will make the strongest contribution to stability, followed by the uncertainty class and with variable response magnitudes having the smallest impact. Our data demonstrate intra-population response diversity may be the norm rather than the exception, and that year-to-year variation in ecological context might mediate, or even reverse, species responses to environmental gradients. This flexibility in the responses of individual species to environmental changes might ensure human-modified environments, like logged and degraded tropical rainforests, are more stable and more resilient ecosystems than expected.

## Acknowledgements

The SAFE Project was supported by the Sime Darby Foundation. Site access and research permits were provided by the Maliau Basin Management Committee, Sabah Foundation, Benta Wawasan, Sabah Softwoods, Innoprise Foundation, Sabah Forestry Department, and the Sabah Biodiversity Centre. RME is supported by the NOMIS Foundation. Data collection was funded by Australian Research Council grant DP140101541; Bat Conservation International; British Council Newton-Ungku Omar Fund; British Ecological Society grant 3256/4035; Cambridge University Commonwealth Fund; Cambridge Trust; Imperial College London SSCP DTP grant NE/L002515/1; Jardine Foundation;

Malaysia Industry Group for High Technology; Panton Trust; Ministry of Education, Youth and Sports of the Czech Republic grant INTER-TRANSFER LTT19018; Primate Society of Great Britain; ProForest; Royal Society of London grant RG130793; Sime Darby Foundation; S.T. Lee Fund; Tim Whitmore Fund; University of Kent; Universiti Malaysia Sabah; UK Natural Environment Research Council grants NE/K016253/1, NE/K016407/1, NE/K016148/1, NE/K0106261/1, NE/K015377/1, NE/L002582/1, NE/P00363X/1, and studentship 1122589; UK Research and Innovation; Universiti Malaysia Sabah, University of Florida Institute of Food and Agricultural Sciences; Varley Gradwell Travelling Fellowship; World Wildlife Fund for Nature. Data collection was supported by Saloni Barsrur, Susan Benedick, Victoria Bignet, Stephen Brooks, Keiron Brown, Stephen Butler, Daniel Carpenter, Kristina Graves, Herry Heroin, Alex Kendall, Darren Mann, Sol Milne, John Mumford, Derek Shapiro, Kathryn Sieving, John Sugau, Elizabeth Telford, Bradley Udell and Bakhtiar Effendy Yahya.

## Author contributions

RME designed the study, conducted the analyses and drafted the manuscript. WDP, CDLO, PA, TdL, GR and CBL supported the data analysis, helped interpret the results and edited the manuscript. All other authors contributed field data and checked the manuscript.

## Materials and methods

Data analysis and construction of figures were conducted in the R v4.02 computing environment (28), using the packages arm (29), dplyr (30), lme4 (31), MuMIn (32), paletteer (33), rstanarm (34, 35), safedata (36) and scales (37).

### Taxa occurrence data

We compiled taxa records from 47 data sources, of which 45 were published on the SAFE Project data repository (38) and the remaining 2 were presented in published papers (39, 40) (Table S1). Data were collected in the eleven-year period 2010 to 2020. We accepted both presence-absence and abundance data for the analysis, but restricted it to records where the sampling locations had known geographic coordinates. Where data sources contained data generated by multiple sampling methods, they were split to consider each method as a separate survey. Similarly, data sources comprising samples collected from multiple years were also split to consider each year as a different survey.

Only identified taxa were analysed, although not all taxa were identified to species. We rejected any taxa identified to less than ordinal level. Morphospecies represent a particularly difficult challenge for analyses of replicability, because the within-survey codes used to identify them are not consistent among surveys. This means taxa cannot be accurately matched and therefore compared among surveys. To surmount this challenge, we aggregated morphospecies by genus within individual surveys, which allowed us to match taxa among surveys. Genus is a commonly used level of taxonomic resolution used in tropical analyses of diverse taxa such as trees (41) and ants (42), and taxa aggregated in this way accounted for just 6 % (n = 32) of all taxa in our analysis.

The median number of surveys per taxon was 3.0 (range: 2-15), and the median number of pairwise comparisons per taxon was also 3.0 (range: 1-105).

### Quantifying forest degradation

Taxa at the SAFE Project were collected along a tropical rainforest degradation gradient that runs from unlogged, old growth forest in strictly protected areas, along a gradient of logging damage going from low-level, selective extraction of individual trees within water catchments through to high-intensity, salvage logged forest with no restrictions on tree harvesting, and ends with oil palm plantation with palms that ranged in age from 5 – 20 years old. We used Aboveground Carbon Density (ACD, Mg.ha^-1^) derived from airborne LiDAR data (43, 44) as a base metric from which to quantify habitat degradation. ACD values ranged from 273 Mg.ha^-1^ in unlogged forest through to just 1 Mg.ha^-1^ in deforested patches. For ease of interpretation, we converted these to a percentage reduction in biomass density relative to the median biomass density observed in unlogged forest (230 Mg.ha^-1^).

We used maps of above-ground carbon density generated from LiDAR data collected in November 2014 (43) and again in April 2016 (44). These dates approximately bracket a salvage logging round during which forest quality was greatly reduced across much of the study area. Each survey was assigned the forest quality metrics that were collected closest in time to the date of the taxa record.

Sampling units varied among the individual surveys, with taxa recorded either at specific point locations (e.g. insect traps), along transects (e.g. fish censes), or within a polygon sampling area (e.g. tree plots). We followed Pfeifer *et al*. (45) by implementing a 1 km buffer area around each sampling unit and averaging the forest quality metrics over all 1-ha pixels within that buffer. We implemented a Gaussian distance weighting that weighted pixels by distance from sampling unit, ensuring pixels located far from the sampling units carried less weight than those immediately adjacent.

We opted to restrict all taxa to be analysed in response to forest quality at a single spatial scale to keep our estimates of response diversity as simple and as conservative as possible. Allowing taxa to vary in the spatial scale of their response among surveys introduces an additional way in which they might have a diversity of response patterns, making it more likely that they would be classified as demonstrating intra-population response diversity. Restricting all taxa to a single spatial scale therefore represents a conservative estimate of the proportion of taxa exhibiting response diversity.

### Quantifying and summarising intra-population response diversity

#### Occurrence models

To combine such disparate sampling methods across such a wide range of taxa, we standardised all taxa records to presence-absence data for analysis. We used univariate, binomial generalised linear models (GLMs) to model taxon occurrence in response to forest degradation, with models fitted to individual surveys. We only fitted models to taxon × survey combinations where the taxon had n ≥ 5 presence records within that particular survey. For each taxon x survey combination, we fitted a model containing a single linear predictor allowing for a sigmoid pattern of occurrence along the forest quality gradient, and estimated the statistical significance of that model using a log-likelihood ratio test comparing the fitted model to a null model.

Controlling for detectability in analyses of species occurrence patterns is often recommended, but such analyses have substantially higher data requirements than typically exist for all taxa within biodiversity surveys (46). In exploratory analyses we found such models routinely failed to converge. This is commonly the case for taxa in the tropics where communities have large numbers of predominantly rare species with low detection probabilities, which are known issues that prevent detectability-based models from converging (46). Moreover, statistical estimates of tropical species responses to ecological gradients do not notably vary between models that incorporate or ignore detection probability (46), so there is no reason to expect the use of detectability-corrected modelling approaches to influence our main conclusions.

#### Categorising response diversity

For the 524 taxa that were detected and modelled from ≥ 2 surveys, we compared model results and grouped taxa according to four classes of response diversity, based on the four classes of ecological context-dependence described by Catford et al (18).

##### A. Invariant

Taxa were considered to have invariant, fully replicable occurrence patterns if all models of that taxa agreed on two criteria:

i. the pattern of significance; binarized into significant (*p* < 0.05) or non-significant (*p* ≥ 0.05); and
ii. the slope and intercept of the fitted model: determined by using the estimate and accompanying standard error to statistically test for a difference among all pairwise combinations of models of the same taxa (47).

##### B. Magnitude

Taxa were considered to vary only in the magnitude of the occurrence pattern if they:

i. had a consistent pattern of significance, but
ii. had slope and/or intercept estimates that varied significantly among surveys.

##### C. Uncertainty

Taxa were considered to vary in their statistical certainty if they had an inconsistent pattern of statistical significance, exhibiting both a statistically significant and statistically insignificant response in one or more surveys respectively.

##### D. Sign

Taxa were considered to vary in sign if they had statistically significant response patterns with opposing signs, meaning in some surveys their occurrence increased with increasing forest degradation but in others their occurrence decreased.

To compare response diversity among broad taxonomic groupings, we categorised all taxa as belonging to one of plant, invertebrate, fish, amphibian, reptile, bird or mammal groups. For each taxonomic group, we took 1,000 bootstrapped samples of the binary responses representing whether the individual taxa within that group lacked (statistical class 1) or exhibited response diversity (one or more of statistical classes 2, 3 or 4). These bootstrapped samples were used to generate distributions describing the proportion of taxa per group with response diversity.

###### Setting a null expectation for the proportion of taxa with response diversity

Statistical techniques, such as the binomial GLM we used to classify response patterns as significant or non-significant, have standard Type I and Type II error rates of 5 and 20 % respectively, meaning the probability of true detection (statistical power) is 80 %. We used these values to set bounds on the proportion of taxa that could have statistically inconsistent results (‘Uncertainty’ response diversity class; Fig. 1C) for spurious reasons. The lower bound is estimated by assuming that all taxa have a “true” response in which they are *not* impacted by habitat degradation. In this scenario, the probability of correctly reporting a non-significant result in one survey (80 %) is multiplied by the probability of incorrectly detecting a significant result where none exists (Type I error; 5 %), meaning 4 % of the pairwise comparisons we analyse would be expected to spuriously report response diversity. Most taxa we examined had > 2 pairwise comparisons, and the probability that at least one pair of surveys for a given taxon returns a spurious result (*P*_*sp*_) scales according to the number of pairwise comparisons (*n*), such that *P*_*sp*(*lower*)_ = 1 − 0.96^*n*^. The upper bound is estimated by assuming that all taxa have a “true” response in which they *are* significantly impacted by the habitat degradation gradient. In this case, we multiply the probability of one survey accurately detecting the significant pattern (power; 80 %) by the probability of the second survey failing to detect the significant pattern that exists (Type II error; 20 %), suggesting 16 % of pairwise comparisons might be expected to spuriously report response diversity. Scaled up to taxon level, this gives a probability of generating a spurious result of *P*_*sp*(*upper*)_ = 1 − 0.84^*n*^.

For each taxon, we estimated *P*_*sp*(*lower*)_ and *P*_*sp*(*upper*)_, and then took the median of each of those two probabilities across all taxa as a null expectation for the proportion of taxa that would exhibit uncertainty response diversity from the spurious accumulation of Type I and Type II errors.

### Sensitivity analyses

#### Minimum number of presence records

We used sensitivity analysis to examine the extent to which our arbitrary threshold of requiring n ≥ 5 presence records per taxon per survey might influence our key results. We progressively restricted our analysis to subsets of taxa that had at n ≥ 5, 10, 15, … 100 presence records within at least two separate surveys, and for each cut-off value calculated the proportion of taxa assigned to each of the four response diversity classes.

#### Critical p-value for statistical significance

Our categorisation of results into uncertainty response diversity (Fig. 1C) is dependent on the *p*-value used to denote statistical significance. We tested the extent to which our results are sensitive to our choice of *p* = 0.05 by repeating the analysis for values of 0.005 ≤ *p* ≤ 0.1 in increments of 0.005.

### Taxonomic bias in the response diversity classes

We used *χ*^2^ goodness of fit tests to determine whether the distribution of taxa within each of the four response diversity classes were random samples of the seven taxonomic groups. We used the proportion of taxa within the full dataset belonging to each of the seven groups as an expected distribution, and assessed the probability that the observed distribution deviated from this. We then used bootstrapping to determine which taxonomic groups were under- or over-represented in each of the four classes. For each class, we took 1000 random draws of the taxa exhibiting that class (sampling with replacement), from which we quantified an expected null distribution of the number of taxa belonging to each of the seven taxonomic groups (Fig. S2). Groups where the observed number of taxa fell below the 2.5 % or above the 97.5 % quantiles of the null distribution were considered to be significantly under- or over-represented respectively.

### Meaningful *versus* spurious intra-population response diversity

We conducted a secondary analysis to classify intra-population response diversity into meaningful or spurious causes. Meaningful response diversity can only be definitively demonstrated when response patterns vary among two surveys that share an exact sampling design and survey method, in which case the variation in response pattern must arise due to unaccounted variation in the environment. To identify cases of confirmed, meaningful response diversity for individual taxa, we first identified all pairs of surveys that contained data on that particular taxon. For each survey pair, we extracted the subset of sampling locations that were present in both surveys and repeated the fitting of binomial GLMs and the categorisation of GLM results into the four response diversity classes. Survey pairs that exhibited one or more of statistical classes B-D (Fig. 1) were confirmed as having meaningful response diversity. This represents a lower bound on the true number of meaningful response diversity cases, as surveys with different spatial designs could also exhibit meaningful response diversity but we are unable to test for it. Taxa that were observed to have response diversity in our main analysis, but for which we could not positively determine the presence of meaningful response diversity in this secondary analysis, were considered to demonstrate spurious response diversity.

We examined three potential causes of meaningful response diversity at our study site: a general year effect; El Niño events that occurred in 2010 and again in 2015/16; and a salvage logging operation that impacted large areas of the landscape over the years 2012-2015. The first of these was tested by quantifying, for each year, the proportion of pairwise comparisons involving that year that exhibited response diversity (Fig. S3A). We quantified this proportion for each of the years for which we had survey data (2010 – 2020 inclusive), and tested for an effect of the number of years separating the two surveys on the whether they had invariant or different response patterns. This was tested using using a binomial GLMM, including taxon identity as a random effect. To test the remaining two hypotheses, we divided pairwise survey comparisons into three groups for each of the two events: (1) *outside*: when each of the individual surveys occurred outside of an El Niño or logging event respectively; (2) *straddling*: when one of the surveys occurred within an El Niño or logging event and the other occurred outside of such events; and (3) *within*: when both surveys occurred within an El Niño or logging event. For each group, we quantified the proportion of pairwise comparisons that exhibited invariant responses (Fig S3B,C). If either of these events were a strong driver of meaningful response diversity, we would expect to see a higher proportion of taxa exhibiting response diversity in survey pairs either straddling or embedded within these disturbance events.

To explore the potential drivers of spurious response diversity, we used generalised linear mixed effects models to examine the effect that researcher-controlled decisions about study design exert on response diversity. We categorised all pairwise combinations of taxon-specific comparisons as exhibiting response diversity or not, and used that as a response variable with taxon identity as a random variable. As fixed effect predictor variables we included the minimum number of occurrences per survey pair (log10-transformed), the minimum number of sample sites per survey pair (log10-transformed), the taxonomic resolution at which that taxa had been identified (species, genus, family or ordinal level), and whether the pair of surveys used the same or different sampling methods. We used minimums rather than the mean or maximum because analyses based on low sample size are more prone to generating spurious and imprecise results, and therefore a pair of studies where one or both have a particularly low sample size are more likely to return inconsistent results. We fitted all variables in a single model and used backwards stepwise model selection using log-likelihood ratio tests to identify significant variables. We calculated the marginal and conditional coefficient of determination (pseudo-r^2^) for the fixed and random effects respectively using the method outlined by Nakagawa and Schielzeth (48).

### Single-survey data in ecology

We reviewed all papers published in *Ecology*, the flagship journal of the Ecological Society of America, in 2021, recording the number of years of empirical data presented in each publication. Out of a total of 263 papers, 246 presented empirical data, of which 100 (41 %) presented data collected from a single-year survey only. A further nine studies (4 %) presented data from multiple datasets, of which one or more was a single-year survey.

### Estimating the probability of detecting a true response

To estimate the probability that we would correctly detect the correct sign of a taxon’s response to the forest degradation gradient, we re-fitted all models using Bayesian hierarchical models in *rstanarm* (34, 35), fitting each taxon with hierarchically-drawn slopes of biomass and intercepts. We used package defaults for model sampling (1000 warm-up iterations and 1000 subsequent sampling iterations across each of 4 chains) with default priors apart from where specified below, and found no evidence of model mis-specification or a lack of convergence (i.e., all 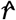 *r*values <1.1 and no divergent transitions in the sampling steps). Specifically, our model was defined as:

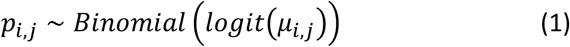

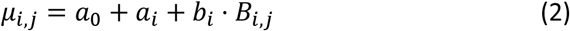

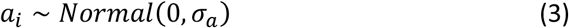

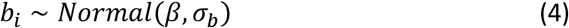

where *p*_i,j_ is the presence/absence (1/0) of a taxon in a given survey (*i*) at a given site (*j*), and *μ*_*i,j*_ is logit-transformed and is proportional to the predicted probability of presence of a species in a given survey (i.e., is a standard Binomial Generalised Linear Model term). *μ*_*i,j*_ itself is then defined by *a*_0_, the overall mean across all years, *a*_*i*_, the contrasts (difference) in the intercept for each survey year, and *b*_*i*_, each survey year’s estimated biomass removal effect (itself multiplied by *B*_*i,j*_, the biomass removal at a given site). The terms *a*_*i*_ and *b*_*i*_ are themselves hierarchically drawn from distributions centred at 0 and *β* and with standard deviations *σ*_*a*_ and *σ*_*b*_, respectively. The Bayesian hierarchical formulation of our approach is central to our method since, for each taxon, it allows us to estimate variation in responses to biomass removal across surveys and also to directly parameterise the distribution of estimated responses in equation 3. From this, we can directly estimate the probability of observing a consistent response across years.

To ensure that our prior specifications were not biasing model results, we repeated our model fits across two extreme prior definitions: ‘correlated’ and ‘variable’ priors. These were fitted across 4 chains, each with 3000 warm-up iterations and 1000 sampling iterations (more samples were needed due to the priors being extreme and therefore slowing model convergence). ‘Correlated’ priors specified regularisation parameters (ζ) of 0.5 for the hierarchical terms, biasing the model such that slopes and intercepts should be more consistent across surveys. ‘Variable’ priors set ζ = 5, such that survey slopes and intercepts were more independent. These two specifications form extremes of a continuum. Our model results were qualitatively identical to those reported in the main text (with ζ set to 1, the default), suggesting that our conclusions are robust to model specification and fitting method.

## SI Appendix Sensitivity analyses

*Minimum number of presence records:* Our results are largely robust to the choice of cut-off value of requiring n ≥ 5 presence records per taxon per survey (Fig. S1A), and our specific cut-off value of n ≥ 5 resulted in the highest proportion of taxa lacking response diversity (having fully invariant responses). This ensures our choice of n ≥ 5 ensures we present results that emphasise the most conservative estimate of response diversity.

*Critical p-value for statistical significance*: Using values of *p* that were lower than 0.05 resulted in a slight reduction in the proportion of taxa demonstrating uncertainty response diversity and a corresponding increase in the proportion of taxa lacking response diversity (Fig. S1B). This effect was most apparent at highly conservative estimates of statistical significance (*p* ≤ 0.02), above which the choice of *p* exerted little influence on our qualitative conclusions.

**Figure S1.**
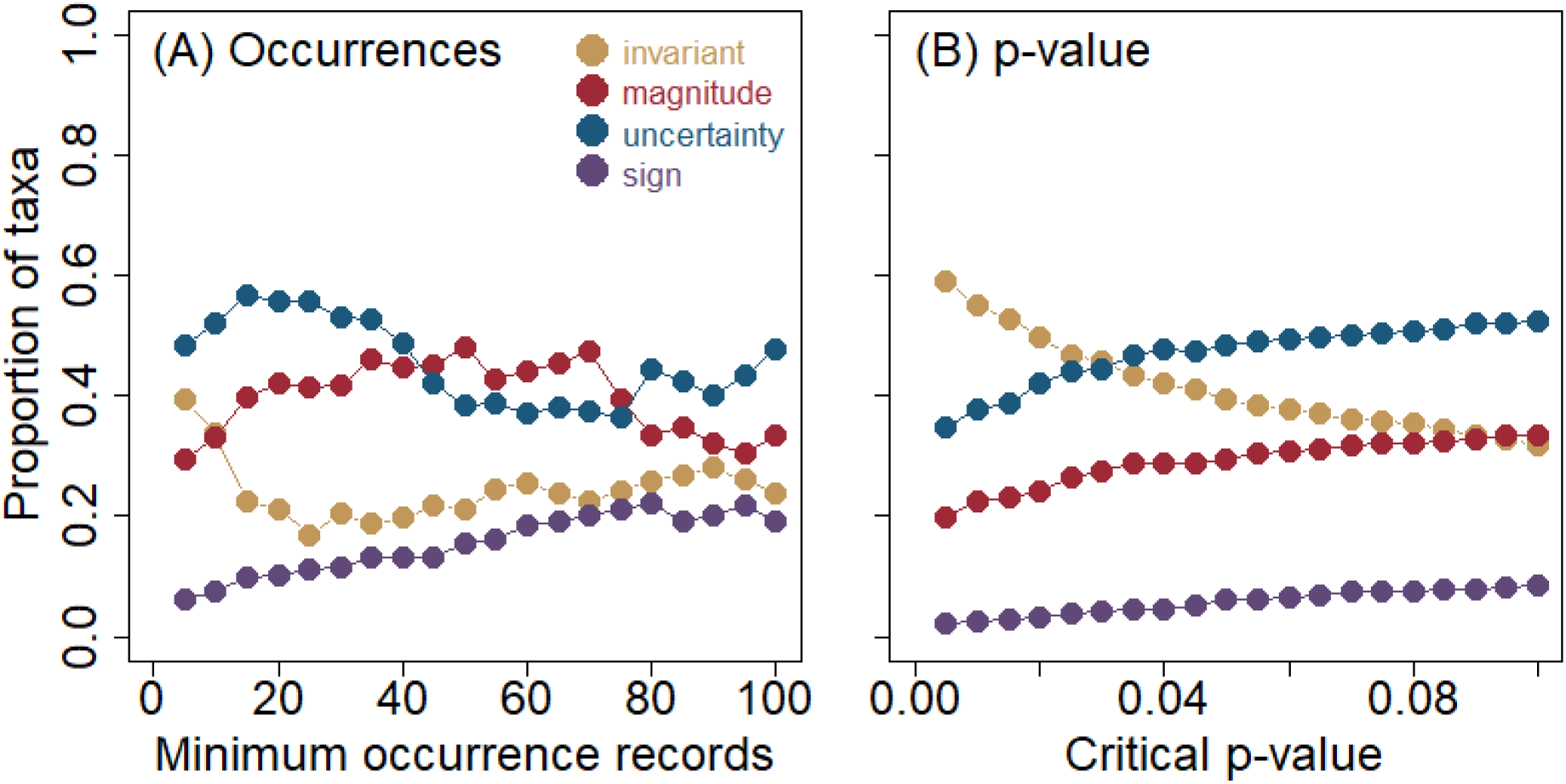
Sensitivity of results to variation in (A) the minimum number of occurrence records required for a taxon to be analysed; and (B) the critical p-value used to denote statistical significance. Values represent the proportion of taxa categorised into each of the four classes of response diversity (Fig. 1).

### Taxonomic bias in the response diversity classes

**Figure S2.**
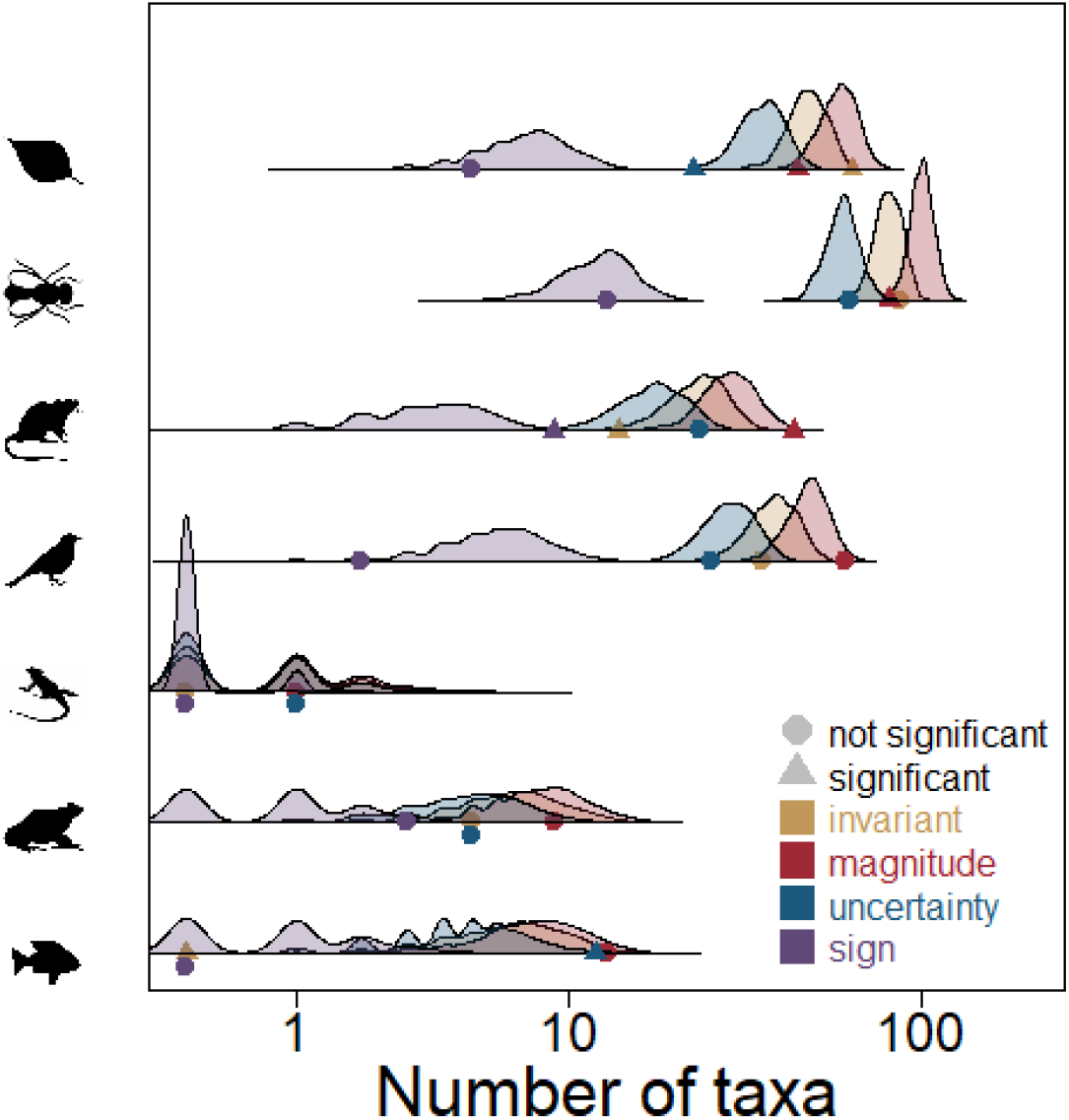
Expected and observed number of taxa within seven broad taxonomic groups exhibiting the four classes of response diversity. Distributions represent, for each taxonomic group and class, a null expectation of the number of taxa that will exhibit that particular class. Points represent the observed number of taxa displaying each class. Significance denotes whether the point sits outside the 95 % quantile of the null distribution.

#### Spurious response diversity

Spurious response diversity is likely driven by methodological differences among surveys of the same taxon in the same landscape, although it’s not clear from our data exactly what researchers should do to eliminate this. The probability that a taxon-specific, pairwise comparison of responses were invariant increased as taxonomic resolution increased (*χ*^2^ = 24.1, df = 3, *p* < 0.001), and there was a paradoxical effect where taxa that were more common were less likely to demonstrate invariant responses (*χ*^2^ = 15.2, df = 1, *p* < 0.001). This likely occurs because rare taxa were less likely to have a significant response to forest degradation (beta regression; *z* = -17.1, *p* < 0.001), and we only found fully invariant responses within generalist taxa that were not impacted by the degradation gradient (e.g. Fig. 1A). Increasing our threshold of n ≥ 5 for inclusion in the analysis further reduced the proportion of taxa with invariant responses (Fig. S1), demonstrating that excluding uncommon species from the analysis would generate even more extreme estimates of response diversity. The probability of responses varying was also increased when pairs of surveys used different sampling methods (*χ*^2^ = 5.25, df = 1, *p* = 0.022), but was not impacted by the minimum sample size of a pair of surveys (*χ*^2^ = 1.67, df = 1, *p* = 0.20). Together, these fixed effects explained just 7 % of the variation in probability of obtaining two invariant surveys, meaning spurious response diversity was not strongly driven by variation in general study design parameters that researchers can exert direct control over. By contrast, the random effect of taxon identity explained 66 % of the variation, indicating the cause of spurious response diversity is almost species-specific. These results indicate that the spurious response diversity we have detected is not obviously generated by high-level design features of studies, and instead appears to be caused by the fine details of individual studies.

**Figure S3.**
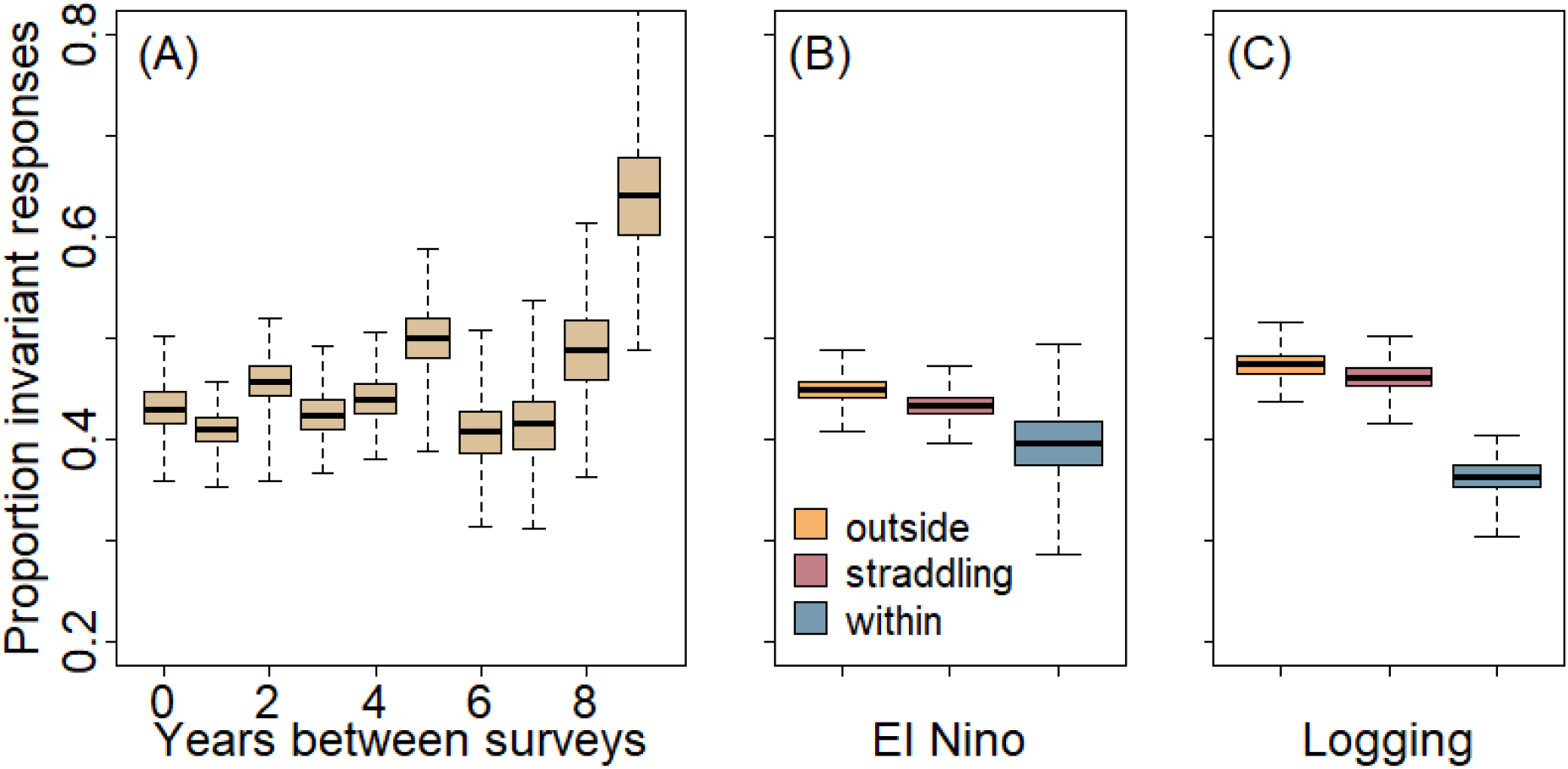
Bootstrapped estimates of the proportion of taxa with invariant response patterns with respect to (A) year of survey, (B) El Niño events and (C) logging events. Thick line represents the median, boxes the 1^st^ and 3^rd^ quantiles, and whiskers the range. In panels (B) and (C), data were categorised into pairwise comparisons of taxon responses to forest degradation in which both surveys were conducted in years during which the event occurred (within), when one survey occurred during an event and the other occurred outside of the event (straddling), and when both surveys occurred outside of the event (outside).

---

**Table S1.** List of data sources compiled for analysis. For each data source, we present: the surname of the first author and a data citation; a weblink to a data publication or, if that is unavailable, then a weblink to a published paper presenting the data; the broad taxonomic grouping(s) that were the focus of the study; the number of taxa that were shared with other surveys; the number of sampling methods used; the number of sampling periods; and the final number of surveys we extracted from that data source.

| First author | Link | Taxon type(s) | No. taxa | No. sampling methods | No. sample periods | No. surveys |
| --- | --- | --- | --- | --- | --- | --- |
| Bernard (49) | https://zenodo.org/record/3908128 | mammal | 21 | 1 | 1 | 1 |
| Bishop (50) | https://zenodo.org/record/1198839 | invertebrate | 22 | 1 | 1 | 1 |
| Both (51) | https://zenodo.org/record/3247631 | plant | 133 | 1 | 1 | 1 |
| Brant (52) | https://zenodo.org/record/1198846 | invertebrate | 21 | 2 | 2 | 3 |
| Carpenter (53) | https://zenodo.org/record/5562260 | invertebrate | 77 | 6 | 1 | 6 |
| Chapman (54) | https://zenodo.org/record/2579792 | mammal | 12 | 1 | 1 | 1 |
| Deere (55) | https://zenodo.org/record/4010757 | mammal | 28 | 1 | 3 | 3 |
| Döbert (56) | https://zenodo.org/record/2536270 | plant | 150 | 1 | 1 | 1 |
| Drinkwater (57) | https://zenodo.org/record/3476542 | invertebrate | 2 | 1 | 2 | 2 |
| Ewers (58) | https://zenodo.org/record/3975973 | invertebrate | 13 | 1 | 1 | 1 |
| Faruk (59) | https://zenodo.org/record/1303010 | amphibian | 2 | 1 | 1 | 1 |
| Fayle (60) | https://zenodo.org/record/3876227 | invertebrate | 22 | 1 | 1 | 1 |
| Fraser (61) | https://zenodo.org/record/3973551 | amphibian | 16 | 1 | 1 | 1 |
| Fraser (62) | https://zenodo.org/record/3981222 | bird | 76 | 1 | 1 | 1 |
| Gray (63) | https://zenodo.org/record/1198302 | invertebrate | 46 | 1 | 1 | 1 |
| Gray (64) | https://zenodo.org/record/3475406 | invertebrate | 4 | 1 | 1 | 1 |
| Gregory (65) | https://zenodo.org/record/3994260 | invertebrate | 1 | 1 | 1 | 1 |
| Hardwick (66) | https://zenodo.org/record/4275386 | invertebrate | 1 | 1 | 1 | 1 |
| Hemprich-Bennett (67) | https://zenodo.org/record/3247465 | mammal | 35 | 1 | 3 | 3 |
| Heon (39) | https://doi.org/10.1890/15-1363 | mammal;<br>bird | 60 | 1 | 6 | 6 |
| Heon (68) | https://zenodo.org/record/3955050 | mammal;<br>bird | 33 | 1 | 7 | 7 |
| Heon (69) | https://zenodo.org/record/1304117 | mammal | 8 | 1 | 1 | 1 |
| Jebrail (70) | https://zenodo.org/record/3475408 | invertebrate | 5 | 1 | 1 | 1 |
| Kendall (71) | https://zenodo.org/record/1237736 | invertebrate | 1 | 1 | 1 | 1 |
| Konopik (72) | https://zenodo.org/record/1995439 | amphibian | 25 | 1 | 3 | 3 |
| Lane Shaw (73) | https://zenodo.org/record/1237732 | invertebrate | 19 | 1 | 1 | 1 |
| Luke (74) | https://zenodo.org/record/5710509 | invertebrate | 7 | 1 | 1 | 2 |
| Luke (75) | https://zenodo.org/record/1198833 | invertebrate | 11 | 2 | 1 | 2 |
| Mackintosh (76) | https://zenodo.org/record/4630980 | invertebrate | 13 | 1 | 1 | 1 |
| Mitchell (40) | https://doi.org/10.1016/j.ecolind.2020.106717 | bird | 132 | 1 | 5 | 5 |
| Mullin (77) | https://zenodo.org/record/3971012 | mammal | 7 | 1 | 2 | 3 |
| Noble (78) | https://zenodo.org/record/3485086 | amphibian | 8 | 1 | 1 | 1 |
| Pianzin (79) | https://zenodo.org/record/3897377 | mammal | 1 | 1 | 1 | 1 |
| Pillay (80) | https://zenodo.org/record/3366104 | bird | 6 | 1 | 2 | 2 |
| Qie (81) | https://zenodo.org/record/3901735 | mammal;<br>bird;<br>invertebrate | 47 | 1 | 1 | 1 |
| Qie (82) | https://zenodo.org/record/1400564 | plant | 275 | 1 | 3 | 3 |
| Seaman (83) | https://zenodo.org/record/5109892 | mammal | 1 | 1 | 1 | 1 |
| Sethi (84) | https://zenodo.org/record/3997172 | bird;<br>amphibian | 227 | 1 | 2 | 3 |
| Shapiro (85) | https://zenodo.org/record/1237720 | invertebrate | 2 | 1 | 1 | 1 |
| Sharp (86) | https://zenodo.org/record/1323504 | invertebrate | 288 | 1 | 3 | 13 |
| Slade (87-89) | https://zenodo.org/record/3247492<br>https://zenodo.org/record/3247494<br>https://zenodo.org/record/3832076 | invertebrate | 71 | 1 | 3 | 3 |
| Slade (90, 91) | https://zenodo.org/record/3906118<br>https://zenodo.org/record/3906441 | invertebrate | 64 | 1 | 2 | 2 |
| Turner (92) | https://zenodo.org/record/5729342 | plant | 93 | 1 | 1 | 1 |
| Twining (93) | https://zenodo.org/record/1237731 | mammal | 2 | 1 | 1 | 1 |
| Vollans (94) | https://zenodo.org/record/3929764 | invertebrate | 1 | 1 | 1 | 1 |
| Wilkinson (95) | https://zenodo.org/record/4072959 | fish | 29 | 3 | 5 | 10 |
| Williamson (96) | https://zenodo.org/record/1487595 | invertebrate | 15 | 1 | 1 | 1 |

## References

1. N. M. Williams et al., Ecological and life-history traits predict bee species responses to environmental disturbances. Biol. Conserv. 143, 2280–2291 (2010).

2. J. P. Suraci et al., Disturbance type and species life history predict mammal responses to humans. Global Change Biol. 27, 3718–3731 (2021).

3. S. R. P.-J. Ross, O. L. Petchey, T. Sasaki, D. W. Armitage, How to measure response diversity. Methods Ecol Evol 14, 1150–1167 (2023).

4. D. Tilman, J. A. Downing, Biodiversity and stability in grasslands. Nature 367, 363–365 (1994).

5. C. Banks-Leite, M. G. Betts, R. M. Ewers, C. D. L. Orme, A. L. Pigot, The macroecology of landscape ecology. Trends Ecol. Evol. 37, 480–487 (2022).

6. C. D. L. Orme et al., Distance to range edge determines sensitivity to deforestation. Nat. Ecol. Evol. 3, 886–891 (2019).

7. J. J. Williams, T. Newbold, Vertebrate responses to human land use are influenced by their proximity to climatic tolerance limits. Divers. Dist. 10.1111/ddi.13282 (2021).

8. M. Tourani et al., Maximum temperatures determine the habitat affiliations of North American mammals. Proc. Natl. Acad. Sci. U.S.A. 120, e2304411120 (2023).

9. K. N. Suding et al., Scaling environmental change through the community-level: a trait-based response-and-effect framework for plants. Global Change Biol. 14, 1–16 (2008).

10. J. H. Hatfield, C. D. L. Orme, J. A. Tobias, C. Banks-Leite, Trait-based indicators of bird species sensitivity to habitat loss are effective within but not across data sets. Ecol. Appl. 28, 28–34 (2018).

11. M. Pacifici et al., Species’ traits influenced their response to recent climate change. Nature Climate Change 7, 205–208 (2017).

12. G. S. Kandlikar, A. R. Kleinhesselink, N. J. B. Kraft, Functional traits predict species responses to environmental variation in a California grassland annual plant community. J. Ecol. 110, 833–844 (2022).

13. J. Leps, F. Bello, P. Šmilauer, J. Doležal, Community trait response to environment: Disentangling species turnover vs intraspecific trait variability effects. Ecography 34, 856–863 (2011).

14. R. M. Ewers et al., A large-scale forest fragmentation experiment: the Stability of Altered Forest Ecosystems Project. Phil. Trans. Roy. Soc. B 366, 3292–3302 (2011).

15. T. Riutta et al., Logging disturbance shifts net primary productivity and its allocation in Bornean tropical forests. Global Change Biol. 24, 2913–2928 (2018).

16. L. N. Joseph, S. A. Field, S. Wilcox, H. P. Possingham, Presence–absence versus abundance data for monitoring threatened species. Conserv. Biol. 20, 1679–1687 (2006).

17. M. Maron, A. Lill, D. M. Watson, R. Mac Nally, Temporal variation in bird assemblages: how representative is a one-year snapshot? Austral Ecol. 30, 383–394 (2005).

18. J. A. Catford, J. R. U. Wilson, P. Pyšek, P. E. Hulme, R. P. Duncan, Addressing context dependence in ecology. Trends Ecol. Evol. 37, 158–170 (2022).

19. M. H. Nunes et al., Recovery of logged forest fragments in a human-modified tropical landscape during the 2015-16 El Niño. Nat. Comm. 12, 1526 (2021).

20. Z. Mehrabi, E. M. Slade, A. Solis, D. J. Mann, The importance of microhabitat for biodiversity sampling. PLoS ONE 9, e114015 (2014).

21. C. Stem, R. Margoluis, N. Salafsky, A. Brown, Monitoring and evaluation in conservation: a review of trends and approaches. Conserv. Biol. 19 (2005).

22. R. Sarremejane et al., Climate-driven hydrological variability determines inter-annual changes in stream invertebrate community assembly. Oikos 127, 1586–1595 (2018).

23. H. Ganjurjav et al., Temperature leads to annual changes of plant community composition in alpine grasslands on the Qinghai-Tibetan Plateau. Environ. Monit. Assess, 190, 585 (2018).

24. S. Cusser, C. Bahlai, S. M. Swinton, G. P. Robertson, N. M. Haddad, Long-term research avoids spurious and misleading trends in sustainability attributes of no-till. Global Change Biol. 26, 3715–3725 (2020).

25. D. G. T. Naidu, S. Roy, S. Bagchi, Loss of grazing by large mammalian herbivores can destabilize the soil carbon pool. Proc. Natl. Acad. Sci. U.S.A. 119, e2211317119 (2022).

26. N. M. Haddad et al., Habitat fragmentation and its lasting impact on Earth’s ecosystems. Sci. Adv. 1, e1500052 (2015).

27. A. E. Noto, T. C. Gouhier, The effects of intraspecific and interspecific diversity on food web stability. Theor. Ecol. 13, 399–407 (2020).

28. R Core Team (2020) R: A language and environment for statistical computing. (R Foundation for Statistical Computing, Vienna, Austria).

29. A. Gelman, S. Yu-Sung (2021) arm: Data analysis using regression and multilevel/hierarchical models. R package version 1.12-2.

30. H. Wickham, R. Francois, L. Henry, K. Muller (2021) dplyr: A grammar of data manipulation.

31. D. Bates, M. Maechler, B. Bolker (2011) lme4: Linear mixed-effects models using S4 classes. R package version 0.999375-39.

32. K. Barton (2020) MuMIn: Multi-Model Inference.

33. E. Hvitfeldt (2021) paletteer: Comprehensive collection of color palettes.

34. B. Goodrich, J. Gabry, I. Ali, S. L. Brilleman (2022) rstanarm: Bayesian applied regression modeling via Stan. R package version 2.21.3.

35. Stan Development Team (2021) RStan: the R interface to Stan. R package version 2.21.3.

36. A. Aldersley, C. D. L. Orme (2019) safedata: Interface to Data from the SAFE Project.

37. H. Wickham, D. P. Seidel (2020) scales: Scale functions for visualization.

38. C. D. L. Orme, R. M. Ewers (2018) The SAFE Project Community.

39. O. R. Wearn, C. Carbone, J. M. Rowcliffe, H. Bernard, R. M. Ewers, Grain-dependent responses of mammalian diversity to land use and the implications for conservation set-aside. Ecol. Appl. 26, 1409–1420 (2016).

40. S. L. Mitchell et al., Spatial replication and habitat context matters for assessments of tropical biodiversity using acoustic indices. Ecol. Indic. 119, 106717 (2020).

41. W. F. Laurance et al., Rapid decay of tree-community composition in Amazonian forest fragments. Proc. Natl. Acad. Sci. U.S.A. 103, 19010–19014 (2006).

42. M. J. W. Boyle et al., Localised climate change defines ant communities in human‐modified tropical landscapes. Func. Ecol. 35, 1094–1108 (2021).

43. T. Jucker et al., Topography shapes the structure, composition and function of tropical forest landscapes. Ecol. Lett. 21, 989–1000 (2018).

44. T. Jucker et al., Estimating aboveground carbon density and its uncertainty in Borneo’s structurally complex tropical forests using airborne laser scanning. Biogeosciences 15, 3811–3830 (2018).

45. M. Pfeifer et al., Creation of forest edges has a global impact on forest vertebrates. Nature 551, 187–191 (2017).

46. C. Banks-Leite et al., Assessing the utility of statistical adjustments for imperfect detection in tropical conservation science. J. Appl. Ecol. 51, 849–859 (2014).

47. C. C. Clogg, E. Petkova, A. Haritou, Statistical methods for comparing regression coefficients between models. Am. J. Sociol. 100, 1261–1293 (1995).

48. S. Nakagawa, H. Schielzeth, A general and simple method for obtaining R2 from generalized linear mixed-effects models. Methods Ecol Evol 4, 133–142 (2013).

49. H. Bernard, K. B. Hee, A. Wong, Importance of riparian reserves and other forest fragments for small mammal diversity in disturbed and converted forest landscapes. https://zenodo.org/record/3908128. (xZenodo, 2020).

50. T. Bishop, R. Ewers, Abundance and morphometrics of ant genera. https://zenodo.org/record/1198839. (xZenodo, 2018).

51. S. Both et al., Functional traits of tree species in old-growth and selectively logged forest. https://zenodo.org/record/3247631. (xZenodo, 2019).

52. H. Brant, J. Mumford, R. Ewers, S. Benedick, Mosquito Data At Safe 2012-2014. https://zenodo.org/record/1198846. (xZenodo, 2018).

53. D. Carpenter et al., The Maliau Quantitative Inventory. https://zenodo.org/record/5562260. (xZenodo, 2021).

54. P. M. Chapman, C. Davison, Small mammals at forest-oil palm edges raw datasets. https://zenodo.org/record/2579792. (xZenodo, 2019).

55. N. J. Deere, Maximizing the value of forest restoration for tropical mammals by detecting three-dimensional habitat associations. https://zenodo.org/record/4010757. (xZenodo, 2020).

56. T. Döbert, B. L. Webber, J. B. Sugau, K. J. M. Dickinson, R. K. Didham, Landuse change and species invasion. https://zenodo.org/record/2536270. (xZenodo, 2019).

57. R. Drinkwater, R. Drinkwater, T. Swinfield, N. J. Deere, Occurrence of blood feeding terrestrial leeches in a degraded forest ecosystem. https://zenodo.org/record/3476542. (xZenodo, 2019).

58. R. M. Ewers, R. Gray, The importance of vertebrates in regulating insect herbivory pressure along a gradient of logging intensity in Sabah, Borneo. https://zenodo.org/record/3975973. (xZenodo, 2020).

59. A. Faruk, Leaf Litter Amphibian Communities. https://zenodo.org/record/1303010. (xZenodo, 2018).

60. T. M. Fayle, K. M. Yusah, R. M. Ewers, M. J. W. Boyle, How does forest conversion and fragmentation affect ant communities and the ecosystem processes that they mediate? https://zenodo.org/record/3876227. (xZenodo, 2020).

61. A. Fraser et al., Amphibian survey of riparian buffer zones at SAFE Project, Borneo. https://zenodo.org/record/3973551. (xZenodo, 2020).

62. A. Fraser, H. Bernard, E. Mackintosh, R. M. Ewers, C. Banks-Leite, Effects of habitat modification on a tritrophic cascade in a lowland tropical rainforest. https://zenodo.org/record/3981222. (xZenodo, 2020).

63. R. Gray, R. Gill, R. Ewers, The Role Of Competition In Structuring Ant Community Composition Across A Tropical Forest Disturbance Gradient. https://zenodo.org/record/1198302. (xZenodo, 2018).

64. R. Gray, E. Slade, A. Chung, O. Lewis, Riparian_Invertebrate_Movement_Data_SAFE. https://zenodo.org/record/3475406. (xZenodo, 2019).

65. N. Gregory, R. M. Ewers, L. Cator, A. Chung, Vectorial capacity of Aedes albopictus across an environmental gradient. https://zenodo.org/record/3994260. (xZenodo, 2020).

66. J. Hardwick et al., The effects of habitat modification on the distribution and feeding ecology of Orthoptera 2015. https://zenodo.org/record/4275386. (xZenodo, 2020).

67. D. Hemprich-Bennett et al., Impacts of rainforest degradation on the diets of the insectivorous bats of Sabah. https://zenodo.org/record/3247465. (xZenodo, 2019).

68. S. Heon, P. M. Chapman, O. R. Wearn, H. Berhard, R. M. Ewers, Core SAFE project small mammal trapping data. https://zenodo.org/record/3955050. (xZenodo, 2020).

69. S. Heon, P. Chapman, H. Bernard, R. M. Ewers, Do Logging Roads Impede Small Mammal Movement In Borneo’S Tropical Rainforests? https://zenodo.org/record/1304117. (xZenodo, 2018).

70. E. W. Jebrail, M. Dahwood, A. H. Fikri, B. Yahya, The effects of progressive land use changes on the distribution, abundance and behavior of vector mosquitoes in Sabah, Malaysia. https://zenodo.org/record/3475408. (xZenodo, 2019).

71. A. Kendall, R. M. Ewers, The Effect Of Forest Modification On Ectoparasite Density And Diversity. https://zenodo.org/record/1237736. (xZenodo, 2018).

72. O. Konopik, Functional diversity of amphibian assemblages along a disturbance gradient. https://zenodo.org/record/1995439. (xZenodo, 2018).

73. I. Lane Shaw, R. M. Ewers, Microclimate Change, Forest Disturbance And Twig-Dwelling Ants. https://zenodo.org/record/1237732. (xZenodo, 2018).

74. S. H. Luke et al., Freshwater invertebrates - diversity and function of stream macroinvertebrates: Effects of habitat conversion and strategies for conservation. https://zenodo.org/record/5710509. (xZenodo, 2021).

75. S. Luke, Ant And Termite Assemblages Along A Tropical Forest Disturbance Gradient In Sabah, Malaysia: A Study Of Co-Variation And Trophic Interactions. https://zenodo.org/record/1198833. (xZenodo, 2018).

76. E. Mackintosh, A. Fraser, C. Banks-Leite, R. M. Ewers, A. Chung, Effect of vertebrate exclusion on ecosystem functioning. https://zenodo.org/record/4630980. (xZenodo, 2021).

77. K. Mullin et al., Bat activity in riparian reserves in forest and oil palm plantations. https://zenodo.org/record/3971012. (xZenodo, 2020).

78. C. Noble, Impacts of habitat disturbance on population health of Bornean frogs. https://zenodo.org/record/3485086. (xZenodo, 2019).

79. A. Pianzin, A. Wong, H. Bernard, M. Struebig, Investigating the distribution and occupancy of otter species across human-modified landscapes in Sabah, Malaysia. https://zenodo.org/record/3897377. (xZenodo, 2020).

80. R. Pillay, R. J. Fletcher, K. E. Sieving, B. J. Udell, H. Bernard, Bioacoustic monitoring reveals shifts in breeding songbird populations and singing behaviour with selective logging in tropical forests. https://zenodo.org/record/3366104. (xZenodo, 2019).

81. L. Qie, E. Telford, R. Nilus, R. Ewers, Increased importance of terrestrial vertebrate seed dispersal in tropical logged forests. https://zenodo.org/record/3901735. (xZenodo, 2020).

82. L. Qie, E. Telford, M. Massam, R. Ewers, Impact Of El Nino Drought On Seedling Dynamics. https://zenodo.org/record/1400564. (xZenodo, 2018).

83. D. Seaman, M. Struebig, H. Bernard, M. Ancrenaz, R. M. Ewers, The effect of tropical forest modification on primate population density and diversity. https://zenodo.org/record/5109892. (xZenodo, 2021).

84. S. Sethi et al., Avifaunal and Herpetofaunal point counts with recorded acoustic data. https://zenodo.org/record/3742834. (xZenodo, 2020).

85. D. Shapiro, R. M. Ewers, Investigating Temperature Tolerance In Mosquito Disease Vectors Across A Land-Use Gradient. https://zenodo.org/record/1237720. (xZenodo, 2018).

86. A. Sharp, M. Barclay, A. Chung, R. Ewers, Beetle Diversity. https://zenodo.org/record/1323504. (xZenodo, 2018).

87. E. M. Slade, E. Bush, D. J. Mann, A. Y. C. Chung, Dung beetle community and dung removal data 2011. https://zenodo.org/record/3247492. (xZenodo, 2019).

88. E. M. Slade, A. Y. C. Chung, J. Parrett, Dung beetle community data 2018. https://zenodo.org/record/3832076. (xZenodo, 2020).

89. E. M. Slade, S. Milne, D. J. Mann, A. Y. C. Chung, J. Parrett, Dung beetle community and dung removal data 2015. https://zenodo.org/record/3247494. (xZenodo, 2019).

90. E. M. Slade, S. Milne, A. Y. C. Chung, J. Williamson, J. Parrett, Dung beetle community and dung removal data 2015. https://zenodo.org/record/3906118. (xZenodo, 2020).

91. E. M. Slade, J. Williamson, A. Y. C. Chung, J. Parrett, H. Heroin, Dung beetle community 2017/18. https://zenodo.org/record/3906441. (xZenodo, 2020).

92. E. C. Turner et al., Tree census data from the SAFE Project 2011-12. https://zenodo.org/record/5729342. (xZenodo, 2021).

93. J. Twining, R. M. Ewers, Terrestrial Scavenger Trapping Data. https://zenodo.org/record/1237731. (xZenodo, 2018).

94. M. Vollans, L. Cator, R. M. Ewers, A. Chung, Investigating the impact of human settlements upon the availability of larval habitats and Aedes albopictus population. https://zenodo.org/record/3929764. (xZenodo, 2020).

95. C. Wilkinson et al., All Fish catch data at the SAFE project 2011-2017. https://zenodo.org/record/3982665. (xZenodo, 2020).

96. J. Williamson, Movement Patterns Of Invertebrates In Tropical Rainforest. https://zenodo.org/record/1487595. (xZenodo, 2018).

